# Viral RNA pUGylation Promotes Antiviral Immunity in *C. elegans*

**DOI:** 10.1101/2025.07.09.663919

**Authors:** David D. Lowe, Aditi Shukla, Scott G. Kennedy

## Abstract

RNA interference (RNAi) is a component of the innate immune systems of many eukaryotes, including *C. elegans*. During RNAi in *C. elegans*, the nucleotidyltransferase RDE-3 modifies the 3’ termini of mRNAs with polyUG (pUG) tails, which recruit RNA-dependent RNA Polymerase (RdRP) enzymes that drive gene silencing by synthesizing antisense small interfering (si)RNAs. During normal growth and development, RDE-3 pUGylates transposon RNAs to silence transposons and protect genomic integrity. How *C. elegans* identifies specific RNAs for pUGylation and whether the pUGylation system is used for other biological purposes is not yet known. Here we show that pUGylation contributes to antiviral immunity in *C. elegans*: During infection of *C. elegans* with Orsay virus, RDE-3 adds pUG tails to viral RNAs, which converts these RNAs into templates for RdRP-based antiviral siRNA production, thereby limiting viral replication. We present evidence that MUT-15 is critical for viral pUGylation because it interacts with RDE-3 and the NYN domain-containing endonuclease RDE-8, thus bridging the enzymes that cleave and pUGylate viral RNA, ensuring efficient antiviral immunity. We conclude that pUGylation promotes antiviral immunity in *C. elegans* and we provide molecular insights into how *C. elegans* identifies and neutralizes its internal and external parasitic threats.

**Importance:** Viruses are a threat to all organisms. Therefore, organisms have evolved numerous systems to recognize and neutralize viruses. Many of these systems, which are referred to as innate immune systems, function by recognizing unique molecular characteristics of viral genetic material. One such innate immune system is RNA interference. RNA interference uses double stranded RNA, which is an obligatory byproduct of replication for many viruses, as a weapon to fight viruses. In this work, we provide molecular insights into how the nematode *C. elegans* uses RNA interference and viral double stranded RNAs to defend itself against viral invaders.

## Introduction

Viruses pose a near-ubiquitous threat to all forms of life, prompting the evolution of diverse innate immune systems that detect and restrict viral replication by recognizing molecular signatures unique to viral nucleic acids. Among these defense mechanisms, RNA interference (RNAi) has emerged as a conserved antiviral strategy in plants and invertebrates (Xie *et al*., 2001; Lu *et al*., 2005; Wilkins *et al*., 2005; Wang *et al*., 2010; Ying *et al*., 2010; Nakahara *et al*., 2012; Guo, Zhang, Wang, Ding, *et al*., 2013; Li *et al*., 2013; Maillard *et al*., 2013, 2019; Tassetto, Kunitomi and Andino, 2017). In *C. elegans*, RNAi is triggered by long double-stranded RNAs (dsRNAs), which are processed by the RNase III-like enzyme DICER into 20–25 nucleotide small interfering RNAs (siRNAs) (Ketting *et al*., 2001; Tabara *et al*., 2002; Pak and Fire, 2007; Guo, Zhang, Wang, Ding, *et al*., 2013). These primary siRNAs are loaded onto the Argonaute protein RDE-1, which guides the complex to complementary target RNAs via Watson-Crick base-pairing (Tabara *et al*., 1999, 2002; Parrish and Fire, 2001; Yigit *et al*., 2006). Target engagement is thought to recruit the endonuclease RDE-8, which cleaves RNAs, enabling RNA-dependent RNA polymerases (RdRPs) to generate secondary siRNAs off RDE-8-cleaved RNAs (Aoki *et al*., 2007; Tsai *et al*., 2015). RdRP-amplified siRNAs are loaded onto secondary Argonautes that reinforce gene silencing (Yigit *et al*., 2006; Vasale *et al*., 2010; Buckley *et al*., 2012). RdRP-based siRNA amplification is a major contributor to the potency and heritability of RNAi in *C. elegans* (Gu *et al*., 2009; Vasale *et al*., 2010; Burton, Burkhart and Kennedy, 2011; Buckley *et al*., 2012; Spracklin *et al*., 2017). Recent discoveries have expanded our understanding of the RNAi pathway by identifying a post-transcriptional RNA modification—poly(UG), or pUG tailing—that facilitates secondary siRNA biogenesis (Preston *et al*., 2019; Shukla *et al*., 2020). The nucleotidyltransferase RDE-3 appends 3’ pUG tails to RNA fragments generated by the coordinated action of RDE-1 and RDE-8 (Chen *et al*., 2005; Shukla *et al*., 2020). pUG tails then adopt G-quadruplex-like structures that recruits RdRPs, enabling efficient secondary siRNA synthesis from pUGylated RNA templates (Roschdi *et al*., 2022). RDE-3 has been shown to pUGylate transposon-derived RNAs during normal development, thereby restricting these endogenous mobile elements and preserving genomic stability (Chen *et al*., 2005; Shukla *et al*., 2020). However, how RDE-3 identifies its RNA substrates, and whether RDE-3 functions with co-factors to achieve selective RNA targeting, remains unclear.

In germ cells, RDE-3 localizes to cytoplasmic foci known as *Mutator* foci (Phillips *et al*., 2012; Shukla *et al*., 2020). The formation of *Mutator* foci requires the low-complexity protein MUT-16, which is thought to scaffold the assembly of small RNA biogenesis machinery within *Mutator* foci (Phillips *et al*., 2012; Uebel *et al*., 2018). Colocalization studies have shown that *Mutator* foci contain several essential RNAi factors, including RDE-3, the exonuclease MUT-7, the RdRP RRF-1, and MUT-15—a protein of unknown domain structure with limited conservation outside nematodes (Phillips *et al*., 2012; Uebel *et al*., 2018; Manage *et al*., 2020). Current models posit that MUT-16 concentrates these components, as well as other factors such as RDE-8, together to facilitate production of the secondary siRNAs required for transposon silencing and germline RNAi (Sijen and Plasterk, 2003; Zhang *et al*., 2011; Phillips *et al*., 2012; Uebel *et al*., 2018). Although many of these proteins are also required for RNAi in somatic tissues, *Mutator* foci have not been detected in the soma (Uebel *et al*., 2018), raising questions about how the pUG RNA and siRNA biogenesis machinery is organized in the absence of *Mutator* foci.

The Orsay virus is a naturally occurring positive-sense single-stranded RNA virus that infects *C. elegans*. Its genome comprises two RNA segments: RNA1 encodes the viral RNA-dependent RNA polymerase, and RNA2 encodes the capsid protein (Félix *et al*., 2011; Jiang, Franz and Wang, 2014; Félix and Wang, 2019). During replication, viral dsRNA intermediates are formed, which are recognized by the RNAi machinery (Wilkins *et al*., 2005; Félix *et al*., 2011; Le Pen *et al*., 2018). For example, RDE-1 and RRF-1 are required to limit Orsay virus replication, establishing RNAi as a functional component of *C. elegans* antiviral immunity (Wilkins *et al*., 2005; Félix *et al*., 2011). Whether RDE-3-mediated pUGylation also contributes to the host’s antiviral response is not known.

Here we show that viral RNA pUGylation is required for Orsay viral immunity in *C. elegans*. We present evidence that MUT-15 promotes viral immunity by recruiting RDE-8, the enzyme responsible for cleaving viral RNAs, into proximity with the pUGylase RDE-3. We also present evidence that the role of MUT-16 in viral immunity is to bring RdRPs into proximity with RDE-3 and RDE-8, thus allowing efficient conversion of pUGylated viral RNA templates into antiviral siRNAs. The findings provide mechanistic insight into how *C. elegans* defends itself against viral pathogens through a unified RNA modification and silencing strategy.

## Results

### RNA pUGylation contributes to innate immunity in *C. elegans*

The NT RDE-3 appends pUG tails to transposon RNAs in the *C. elegans* germline to limit transposon replication (Shukla *et al*., 2020). We wondered if the pUG system might play additional roles in protecting *C. elegans* against nucleic acid parasites. To test this idea, we infected wild-type (WT), animals harboring an *rde-3* deletion that removes conserved domains (RDE-3(-)), animals harboring two different RDE-3 catalytic site mutations (G366R or D189N), or RDE-3(G366R) animals reverted to wild-type (RDE-3(GR366G-reverted)) with Orsay virus and used qRT-PCR to monitor levels of Orsay viral RNAs (ORV1 and ORV2) in these animals four days post-infection. We observed a ∼6-7 log2-fold increase in viral ORV1 and ORV2 RNA (henceforth, viral load) in animals that lacked functional RDE-3, indicating that RDE-3 is needed to limit Orsay viral replication in *C. elegans* (Figure 1A/B).

**Figure 1.**
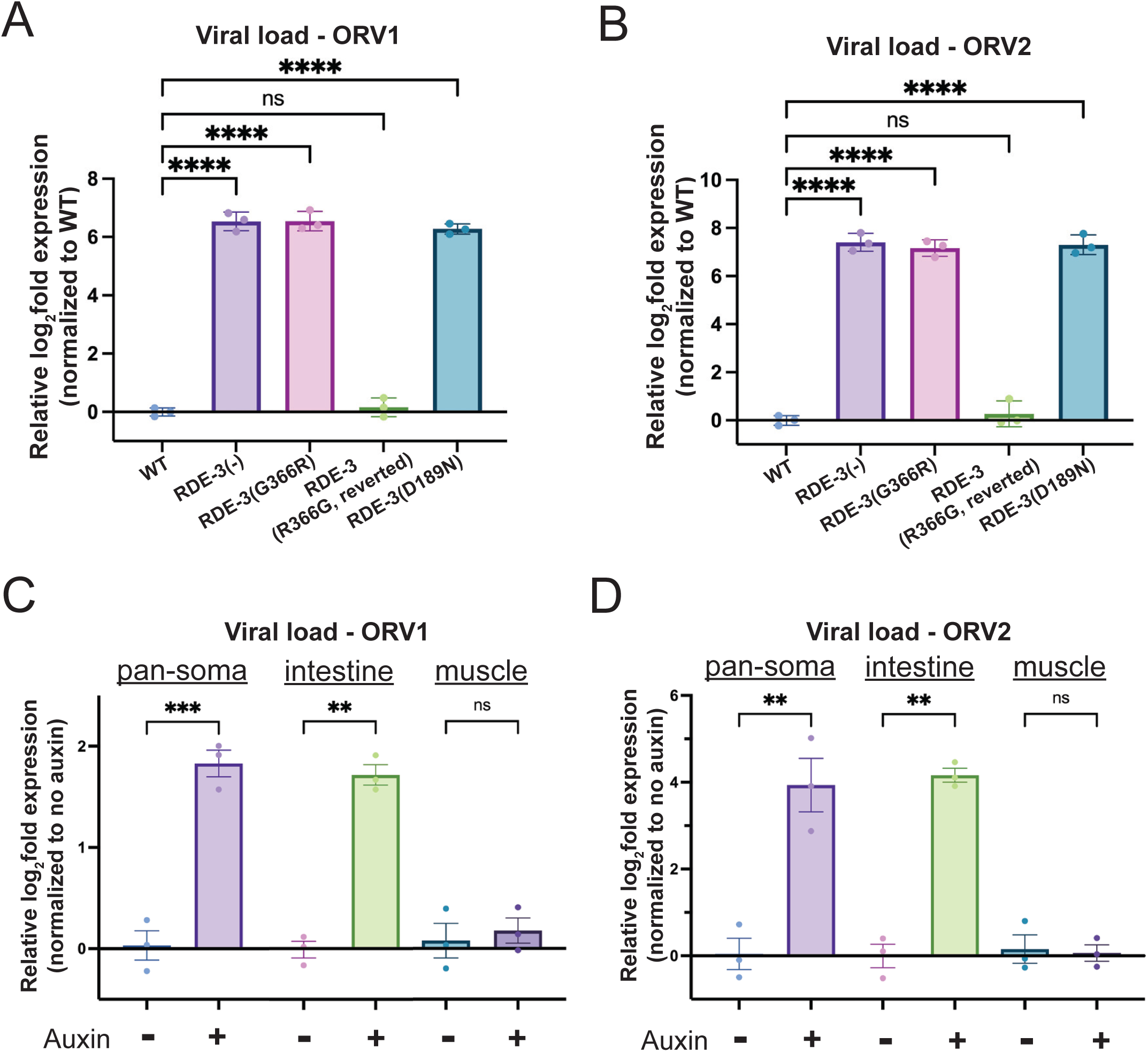
RDE-3 limits viral replication in intestinal cells. **(A-B)** qRT-PCR to determine viral load in WT, *rde-3(ne3370)* [RDE-3(-)], *rde-3(ne298)* [RDE-3(G366R)], *rde-3(gg679)* [RDE-3(R366G) reversion], and *rde-3(r459)* [RDE-3(D189N)] animals detecting **(A)** ORV RNA1 (ORV1) and **(B)** ORV RNA2 (ORV2). Data are normalized to housekeeping genes *eft-2* and *cdc-42* and viral load in WT-no infection was defined as zero. Unpaired, one-way ANOVA. n=3. **(C-D)** qRT-PCR to determine viral load, ORV1 **(C)** and ORV2 **(D)** relative to the genome-encoded UG-containing mRNA *gsa-1* and normalized to non-auxin treated animals. Paired, two-tailed t-test. n=3 ns - not significant,* p<0.05, ** p<0.01, *** p<0.001, **** p<0.0001.

Orsay virus infects and replicates within the intestinal cells of *C. elegans* (Félix *et al*., 2011; Franz *et al*., 2014; Guo *et al*., 2020). To determine the site of RDE-3’s antiviral activity, we employed the auxin-inducible degron (AID) system to deplete RDE-3 in a tissue-specific manner (Zhang *et al*., 2015; Ashley *et al*., 2021). We generated animals that expressed GFP::AID::RDE-3 in all tissues (Shukla *et al*., 2020) and the F-box Transport Inhibitor Response 1 (TIR1), which is needed for auxin-induced degradation, in specific tissues (Zhang *et al*., 2015; Ashley *et al*., 2021). Pan-soma or intestinal depletion, but not muscle depletion, of RDE-3 led to animals unresponsive to RNAi targeting the intestinally expressed *dpy-6* mRNA, which indicates tissue-specific RDE-3 depletion was successful (Figure S1). We performed auxin-induced degradation of RDE-3 in all somatic cells (pan-soma) or in muscle or intestinal cells of animals infected with Orsay virus. Pan-soma or intestinal depletion, but not muscle depletion, of RDE-3 resulted in ∼2-4 log2-fold increase in viral load post-Orsay virus infection (Figure 1C/D). The increase in viral load observed after intestinal RDE-3 depletion indicates RDE-3 acts cell-autonomously to limit viral replication. The failure of Orsay virus to accumulate to the same level exhibited by *rde-3(-)* animals after intestinal RDE-3 knockdown may be due to incomplete knockdown of RDE-3 in these experiments, or it may indicate additional, non-autonomous roles for RDE-3 in antiviral immunity.

Because the pUGylase RDE-3 restricts viral RNA expression, we hypothesized that pUGylated viral RNAs might be produced during Orsay virus infection. To test this hypothesis, we performed RT-PCR specific for pUG-modified RNAs to detect pUGylated OrV RNA1 transcripts in wild-type (WT) or RDE-3(-) animals that had been infected with Orsay. The analyses revealed Orsay virus-derived pUGylated RNAs generated post-infection, which required RDE-3 for their biogenesis (Figures 2A/B). An *in vitro* synthesized *gfp* pUG RNA, which was spiked-in during library preparation, was detected at similar levels in both WT and RDE-3(-) animals, indicating failure to detect Orsay pUGs in RDE-3(-) animals was not due to technical issues (Figure S2). Additionally, we employed Nanopore-based cDNA sequencing to sequence pUGylated RNAs in RNA isolated from WT and RDE-3(-) animals infected with Orsay virus (Methods). Nanopore sequencing detected ORV1 and ORV2 RNAs possessing 3’ non-templated pUG tails in WT animals, but not in RDE-3(-) animals (Figure 2C). Analysis of the Nanopore sequencing data revealed that pUG tails were added to fragments of ORV1 and ORV2 RNAs and that sites of pUGylation on these RNAs were distributed across both viral RNAs (Figure 2C). Collectively, these findings show that Orsay RNA undergoes RDE-3-dependent pUGylation.

**Figure 2.**
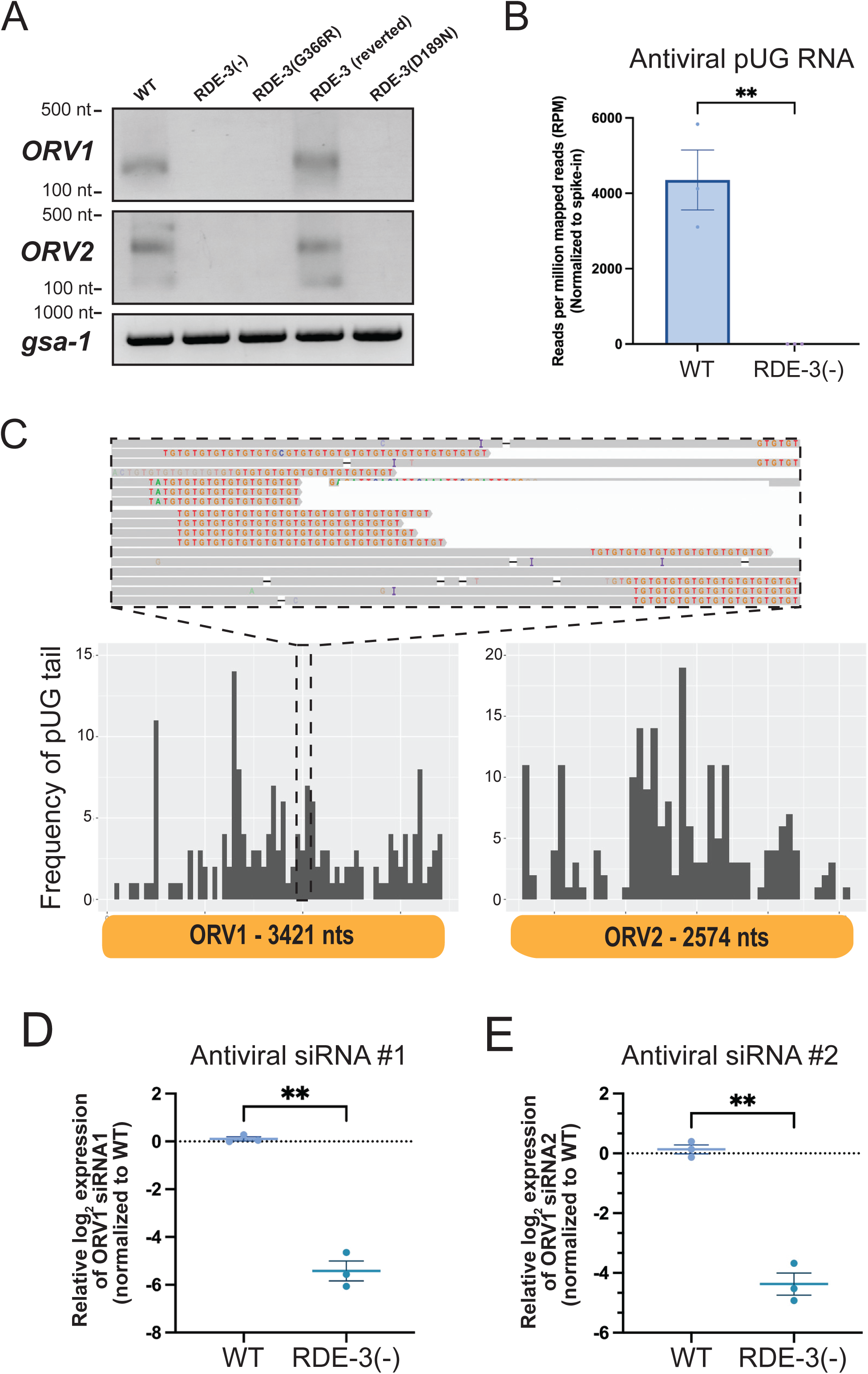
Orsay viral RNAs are pUGylated in an RDE-3 dependent manner. **(A)** RT-PCR detecting ORV1 and ORV2 pUG RNAs, or the control pUG-containing mRNA *gsa-1*, in animals of the genotypes described in Figure 1. **(B)** Number of viral pUG RNAs from WT and *rde-3(ne3370)* [RDE-3(-)] infected with Orsay virus as identified by Nanopore-based pUG RNA sequencing. Data were normalized to the number of sequenced GFP::pUG RNAs, which were spiked into RNA preparations prior to library construction (see Figure S2). Unpaired, two-tailed t-test. n=3. **(C)** (Top) IGV screenshot of ORV1 and ORV2 pUG RNAs sequenced on the Nanopore platform. Gray indicates reads mapped to ORV1 or ORV2 and pUG tails are shown as TG repeats (red and orange). (Bottom) Histogram showing a representative WT replicate with numbers of pUG tails (binned in 50 nucleotide increments) identified across the viral genome: ORV1 (left) and ORV2 (right). Similar results were observed in two other replicates. **(D-E)** Taqman assays on RNA isolated from WT and *rde-3(ne3370)* [RDE-3(-)] animals infected with Orsay virus to detect antiviral siRNA #1 **(D)** or siRNA #2 **(E)** relative to a control snRNA, *U18*. Paired, two-tailed t-test. ns - not significant,* p<0.05, ** p<0.01, *** p<0.001, **** p<0.0001.

Following infection, *rde-3* mutants exhibited elevated viral loads and lack detectable viral pUG RNAs, suggesting that pUGylated viral RNAs contribute to antiviral immunity. How might pUGylated viral RNAs restrict viral replication? RDE-3 generates RNAs with pUG tails that form atypical G-quadruplex RNA structures, which interact with the RNA-dependent RNA polymerases (RdRPs) RRF-1 and EGO-1 (hereafter referred to collectively as RdRP) (Shukla *et al*., 2020; Roschdi *et al*., 2022). RdRP utilizes pUGylated RNAs as templates to synthesize antisense siRNAs, which function in *trans* to silence complementary RNAs (Shukla *et al*., 2020; Roschdi *et al*., 2022). Thus, viral pUG RNAs could facilitate antiviral immunity by promoting RdRP-mediated antiviral siRNA synthesis. Supporting this model, previous work demonstrates increased Orsay viral loads in animals deficient for RdRP (Ashe *et al*., 2013; Le Pen *et al*., 2018; Long, Meng and Lu, 2018). Using custom-designed Taqman probes, we quantified levels of two antiviral siRNAs, which were identified in previous studies (Ashe *et al*., 2013; Le Pen *et al*., 2018; Long, Meng and Lu, 2018), in WT or RDE-3(-) animals infected with Orsay. The analysis showed that RDE-3(-) animals produced approximately ∼4–6 log2-fold fewer antiviral siRNAs than WT animals after infection (Figure 2D/E). These results suggest that viral pUGylated RNAs inhibit viral replication by acting as templates for antiviral siRNA biogenesis.

### *In silico* predictions of RDE-3 interactors

To explore the molecular basis of RDE-3-based RNA pUGylation, and to identify potential RDE-3 co-factors, we leveraged immunoprecipitation-mass spectrometry (IP-MS) data that previously identified 21 proteins enriched in MUT-16 pulldowns, including the pUGylase RDE-3 (Tsai *et al*., 2015; Uebel *et al*., 2018; Manage *et al*., 2020). To predict potential RDE-3 interacting proteins, we used AlphaFold2 to assess whether any MUT-16–interacting proteins might be predicted to interact directly with RDE-3. As a specificity control, we included 100 randomly selected, conserved, germline-expressed *C. elegans* proteins in the analysis. We evaluated two AlphaFold2 output metrics to predict protein-protein interactions: (1) interface-predicted template modeling (ipTM) scores, which estimate interaction likelihood (Jumper *et al*., 2021), and (2) the number of independent structure predictions (out of five) that support the interaction, which indicates prediction robustness (Lim *et al*., 2023). This analysis identified two high-confidence RDE-3 interactors: MUT-15 and UAF-1 (Figure 3A). Given prior evidence that RDE-3 and MUT-15 co-localize in germ cells and are both required for RNA interference and transposon silencing (Chen *et al*., 2005; Zhang *et al*., 2011; Phillips *et al*., 2012; Uebel *et al*., 2018; Shukla *et al*., 2020), we focused further investigation on MUT-15. To assess specificity of the predicted MUT-15-RDE-3 interaction, we used AlphaFold2 to test for predicted interactions between MUT-15 and 40 other nucleotidyltransferases (NTs) encoded in the *C. elegans* genome. RDE-3 was the only NT predicted to interact with MUT-15, indicating a degree of specificity in the Alphafold predicted MUT-15–RDE-3 interaction (Figure 3A). A predicted alignment error (PAE) plot for the predicted MUT-15-RDE-3 interaction is shown in Figure 3B. A three dimensional (3D) representation, and four other independent PAE predictions, of the predicted MUT-15-RDE-3 complex are shown in Figure S3A.

**Figure 3.**
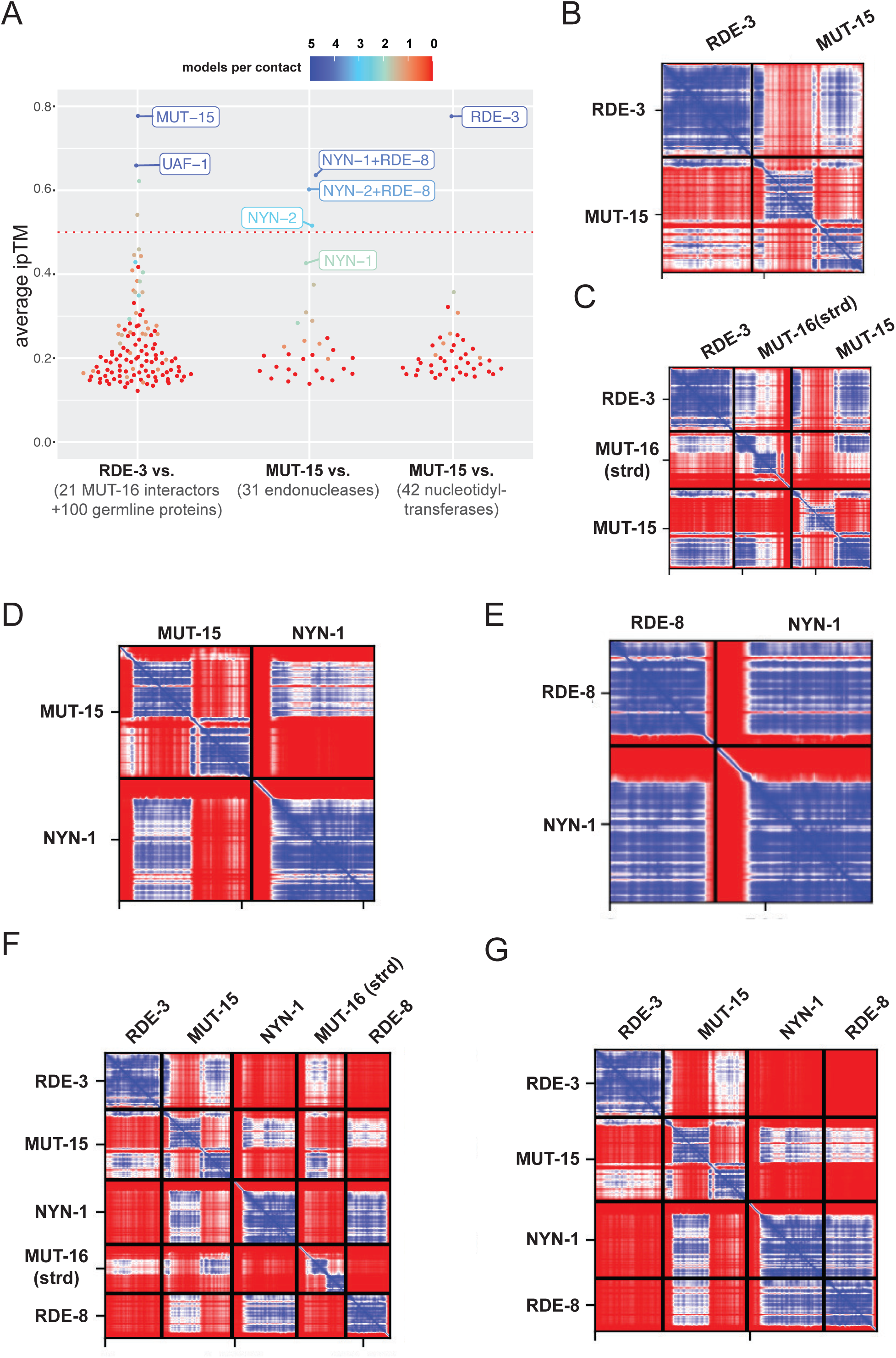
*In silico* prediction of RDE-3 interacting proteins. **(A)** Summary of Alphafold2 screens showing average ipTM scores for all predictions. Average number of models indicating an interaction per predicted contact is indicated by color: Color scale is shown at the top. (Left) RDE-3 queried against 100 conserved germline proteins and 21 known MUT-16 interacting proteins. (Middle) MUT-15 queried against 42 *C. elegans* nucleotidyltransferases. (Right) MUT-15 queried against 29 endonucleases and RDE-8:NYN-1 or RDE-8:NYN-2. **(B-G)** Top ranked PAE plots for the indicated Alphafold2 protein-protein interaction predictions are shown. PAE plots are color-coded to show predicted distances between residues: 0 Angstroms (blue) to 30 or greater Angstroms (red).

Given that RDE-3 and MUT-15 co precipitate with MUT-16, we asked if Alphafold2-Multimer might predict a direct interaction between either RDE-3 and MUT-16 or MUT-15 and MUT-16. Previous studies have demonstrated that co-localization of RDE-3 and MUT-15 with MUT-16 in germ cells depends upon amino acids 89–384 of MUT-16, which is the only segment of MUT-16 predicted to adopt secondary structure (hereafter MUT-16(strd)) (Phillips *et al*., 2012; Uebel *et al*., 2018). AlphaFold2-Multimer predicted that MUT-15, but not RDE-3, interacted with MUT-16(strd). PAE plots of the predicted MUT-15-MUT-16 interaction are shown in Figure S3B. Modeling of RDE-3, MUT-15, and MUT-16(strd) together resulted in the prediction of a trimeric complex, in which MUT-15 bridges RDE-3 and MUT-16(strd). PAE plots and a 3D representation of the predicted MUT-15-RDE-3-MUT-16(strd) interaction are shown in Figure 3C and Figure S3C, respectively.

RDE-8, NYN-1, and NYN-2 are NYN (<u>N</u>edd4-BP1, <u>Y</u>acP <u>N</u>ucleases) domain-containing endonucleases. RDE-8 is required for RNAi and NYN-1 and NYN-2 are redundantly required for RNAi in *C. elegans* (Tsai *et al*., 2015). During RNAi, RDE-8 is thought to endonucleolytically cleave mRNAs targeted for silencing by dsRNAs and the Argonauts RDE-1 (Tsai *et al*., 2015). NYN-1 and NYN-2 lack catalytic residues required for endonuclease activity (Tsai *et al*., 2015), suggesting that NYN-1 and NYN-2 play redundant, non-catalytic roles in RNAi. MUT-16 co-precipitates with RDE-8, NYN-1, and NYN-2 (Tsai *et al*., 2015) and MUT-16(strd) is required for RDE-8, NYN-1, and NYN-2 to colocalize with MUT-16 in *C. elegans* germ cells (Uebel *et al*., 2018). Thus, RDE-8, NYN-1, and NYN-2 are strong candidates for proteins that could interact with MUT-15, RDE-3, or MUT-16. To probe potential interactions between RDE-8, NYN-1, or NYN-2 with MUT-15, RDE-3, or MUT-16 we again utilized AlphaFold2-Multimer modeling. The resulting predictions showed that NYN-1 and NYN-2, but not RDE-8, were predicted to interact with MUT-15 through the same interface on MUT-15, suggesting a mutually exclusive interaction between NYN-1 or NYN-2 with MUT-15. The predicted interaction surface on MUT-15 for NYN-1/2 was distinct from the predicted RDE-3 interaction surface on MUT-15. The predicted mutually exclusive interactions of NYN-1 or NYN-2 with MUT-15 are consistent with the redundant role played by NYN-1 and NYN-2 in RNAi. PAE plots of the predicted NYN-1-MUT-15 and NYN-2-MUT-15 interactions are shown in Figure 3D and Figure S4A, respectively. To assess specificity of the predicted NYN-1/2-MUT-15 interaction, we examined 26 additional *C. elegans* endonucleases for predicted binding to MUT-15. Only NYN-1 and NYN-2 displayed predicted interactions, indicating a degree of specificity in the NYN-1/2-MUT-15 predictions (Figure 3A). Although a direct interaction between RDE-8 and MUT-15 was not predicted, additional modeling with just RDE-8 and NYN-1/2 revealed predicted interactions between RDE-8 and NYN-1 or NYN-2, which is consistent with previous *in vivo* studies (Tsai *et al*., 2015). PAE plots and 3D representations of the predicted NYN-1/2-RDE-8 interactions are shown in Figure 3E and Figure S4B, respectively.

When RDE-8, NYN-1 or NYN-2, MUT-15, MUT-16(strd), and RDE-3 were modeled simultaneously, Alphafold2-Multimer predicted a heteropentameric complex. In this predicted complex, MUT-15 bridges the RNA pUGylase RDE-3, MUT-16, and NYN-1/2, while NYN-1/2 connects the endonuclease RDE-8 to the other components of the predicted complex (Figure 3E and Figure S5). When MUT-16 was not included in the prediction, the other four proteins (RDE-8, NYN-1 or NYN-2, MUT-15, and RDE-3) were predicted to form a similar complex, which lacked MUT-16 (Figure 3F and Figure S6). PAE plots and 3D representation of the predicted heteropentameric (with MUT-16) and heterotetrameric complex (without MUT-16) are shown in Figure 3E/F and Figure S5-6. Neither RDE-3 nor MUT-16 were predicted to interact with NYN-1, NYN-2, or RDE-8 within these complexes. Because data presented below will show that RDE-3, MUT-15, NYN-1/2, and RDE-8, but not MUT-16, are required for viral RNA pUGylation, we refer to the four-protein heterotetrameric complex containing RDE-3, MUT-15, NYN-1/2, and RDE-8 as the predicted pUGasome.

### Co-immunoprecipitation studies support predicted MUT-15 interactions and the general architecture of the predicted pUGasome

To experimentally test the AlphaFold2-Multimer-based predictions of a pUGasome complex, we introduced epitope tags at the endogenous loci of the four core predicted components: 3XFLAG::RDE-3, MUT-15::3XHA, 3XV5::RDE-8, and 3XV5::NYN-1/NYN-2. Prior studies have established the necessity of RDE-3, RDE-8, and MUT-15 for efficient RNA interference (RNAi) in *C. elegans*. To confirm that the introduced tags did not compromise protein function, we performed RNAi assays targeting *dpy-6*, a gene whose silencing results in a characteristic Dumpy (*Dpy*) phenotype in wild-type animals. Consistent with expectations, animals harboring null mutations in *rde-3*, *mut-15*, or *rde-8* failed to exhibit the Dpy phenotype upon exposure to *dpy-6* RNAi (Figure S7A). When animals expressing 3X FLAG::RDE-3, MUT-15::3X HA, or 3X V5::RDE-8 were exposed to *dpy-6* RNAi they became dumpy (Figure S7B), indicating that the introduced tags did not disrupt protein function.

We conducted four Co-IP experiments to test AlphaFold2-predicted interactions within the pUGasome. First, we examined the predicted interaction between RDE-3 and MUT-15 by conducting co-immunoprecipitation (co-IP) assays, which confirmed an *in vivo* physical interaction between RDE-3 and MUT-15 (Figure 4A). Second, AlphaFold2 modeling suggested residues 2–44 of MUT-15 are essential for interaction with RDE-3 (Figure 4B). Modeling a truncated MUT-15 variant lacking residues 2–44 (henceforth referred as MUT-15(Δ2–44)) failed to predict binding with RDE-3 *in silico* (Figure S8). To assess if a.a. 2-44 of MUT-15 were required for MUT-15’s interaction with RDE-3 *in vivo*, we generated a *mut-15(Δ2–44)* allele via CRISPR-Cas9 genome editing. MUT-15(Δ2–44) animals were unresponsive to *dpy-6* RNAi, demonstrating a functional requirement for residues 2–44 of MUT-15 in RNAi (Figure 4C). Additionally, while MUT-15(Δ2–44) was expressed, it failed to co-immunoprecipitate with RDE-3 (Figure 4D). Third, we tested whether the predicted RDE-3–MUT-15 interaction requires MUT-16: AlphaFold2 predicts an interaction that is independent of MUT-16. Co-IP experiments in MUT-16(-) animals showed that RDE-3 and MUT-15 co-precipitated in the absence of MUT-16, validating a MUT-16-independent interaction (Figure 4A). Fourth, we assessed whether RDE-3 interacts with RDE-8 or NYN-1/2 through MUT-15, as computationally predicted. Indeed, RDE-3 co-immunoprecipitated with RDE-8 and NYN-1 in wild-type animals, but not in MUT-15(-) animals, consistent with the idea that MUT-15 bridges RDE-8 and NYN-1 to RDE-3 (Figure 4E). Together, the co-IP analyses support the general architecture of the pUGasome, as predicted by Alphafold2.

**Figure 4.**
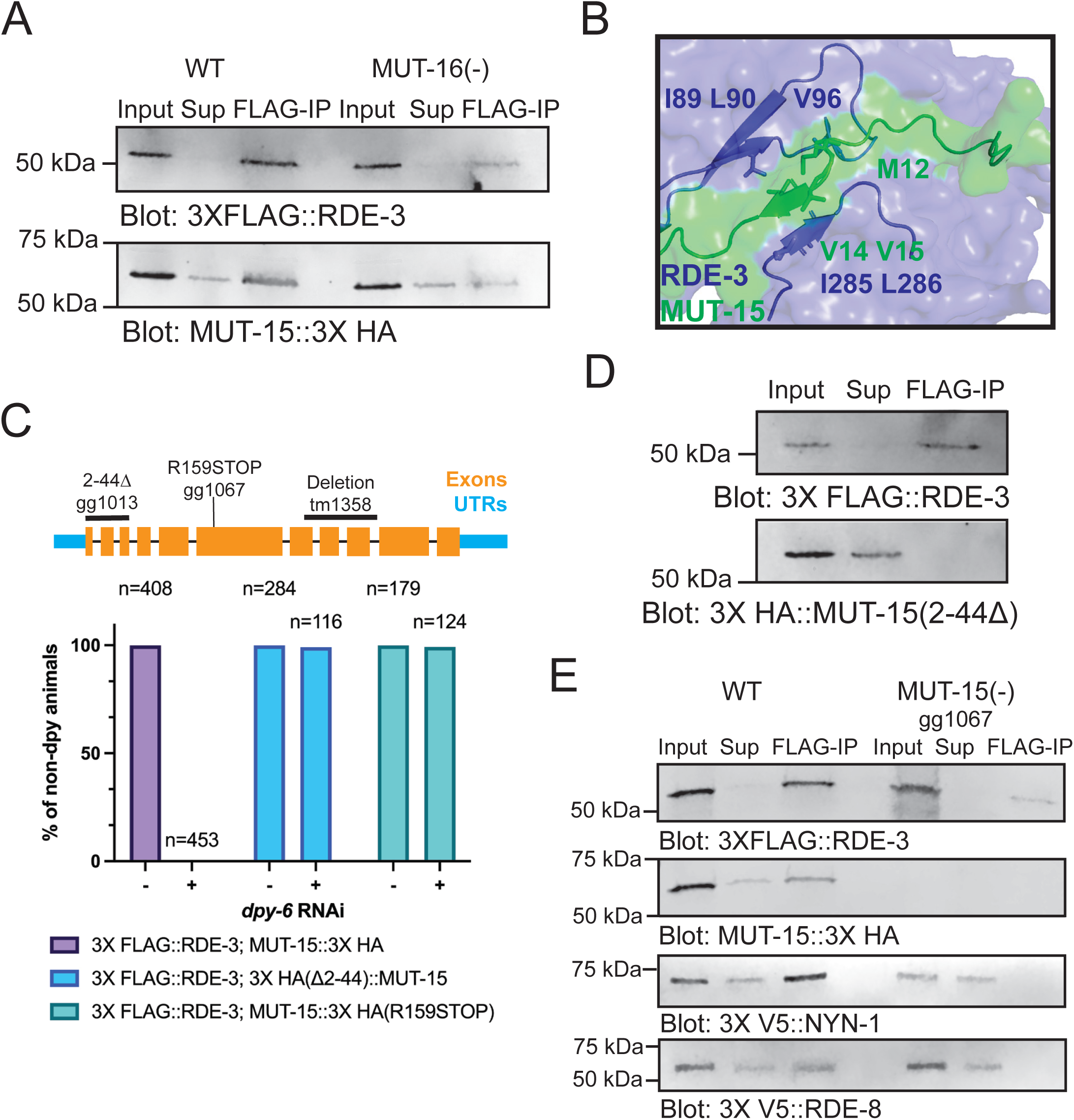
*In vivo* tests of Alphafold2 predictions. **(A)** 3X FLAG::RDE-3 was immunoprecipitated from extracts with ɑ-FLAG antibodies. *mut-16(pk710)* background is indicated as MUT-16(-). 10% of input, 2% of supernatant (Sup), or 10% of FLAG-IPs were subjected to Western blotting with FLAG or HA antibodies to detect 3XFLAG::RDE-3 or MUT-15::3XHA. **(B)** 3D depiction of Alphafold2 model for a.a. 1-50 of MUT-15 (green) interacting with the surface of RDE-3 (blue). **(C)** (Top) *mut-15* gene structure, including mutant alleles used in this study. (Bottom) RNAi assay showing percentage of non-dumpy animals after WT, *mut-15(gg1013)* [MUT-15(Δ2-44)], and *mut-15(gg1067)* [MUT-15(R159STOP)] animals were subjected to *dpy-6* RNAi. **(D)** FLAG-immunoprecipitation followed by FLAG or HA blotting to detect 3XFLAG::RDE-3 and 3XHA::MUT-15(Δ2-44). 8% of input, 4% of supernatant (Sup), and 8% of FLAG-IP were probed. **(E)** FLAG-immunoprecipitation followed by blotting with indicated antibodies for 3XFLAG::RDE-3, MUT-15::3XHA, 3XV5::RDE-8, or 3XV5::NYN-1 in WT or *mut-15(gg1067)* [MUT-15(-)] animals. 10% of input, 2% of supernatant (Sup), and 10% of FLAG-IP were probed.

### Components of pUGasome are required for viral pUG RNA synthesis and antiviral immunity

To investigate the *in vivo* significance of the predicted pUGasome, we examined the effects of disrupting each component on viral replication, antiviral pUG RNA synthesis, and antiviral siRNA synthesis. We first quantified Orsay virus RNA levels via qRT-PCR in wild-type animals and mutants deficient in individual pUGasome components: RDE-3(-), RDE-8(-), NYN-1(-); NYN-2(-), MUT-15(-), or MUT-15(Δ2–44). All five mutant strains exhibited significantly increased viral RNA loads, approximately ∼4–5 log2-fold higher than wild-type controls (Figure 5A/B and Figure S9A). These results demonstrate that each pUGasome component, including amino acids 2–44 of MUT-15, which are required for MUT-15 to interact with RDE-3, are required for suppressing viral replication. These observations align with previously reported findings for *rde-8* mutants (Tsai *et al*., 2015). To determine if the predicted components of the pUGasome are required for viral RNA pUGylation, we measured ORV1 pUG-modified RNAs by qRT-PCR in animals lacking components of the predicted pUGasome. Animals lacking any of the four predicted pUGasome components failed to produce detectable ORV1 pUG RNAs after infection (Figure 5C), establishing an essential role for the predicted pUGasome components in mediating viral RNA pUGylation. We next quantified antiviral secondary siRNAs generated in response to Orsay virus infection using Taqman-based assays. Absence of any pUGasome component resulted in loss of antiviral siRNA production (Figure 5D and Figure S9B). Collectively, the results establish a critical role for the four predicted components of the pUGasome —RDE-3, MUT-15, RDE-8, and NYN-1/2—in three aspects of *C. elegans* antiviral defense: (1) pUGylation of viral RNAs (henceforth, antiviral pUG RNAs), (2) synthesis of antiviral siRNAs, and (3) suppression of viral replication. The results support the existence of a functional pUGasome and underscore the importance of this predicted pUGasome in orchestrating RNA-based innate immune responses.

**Figure 5.**
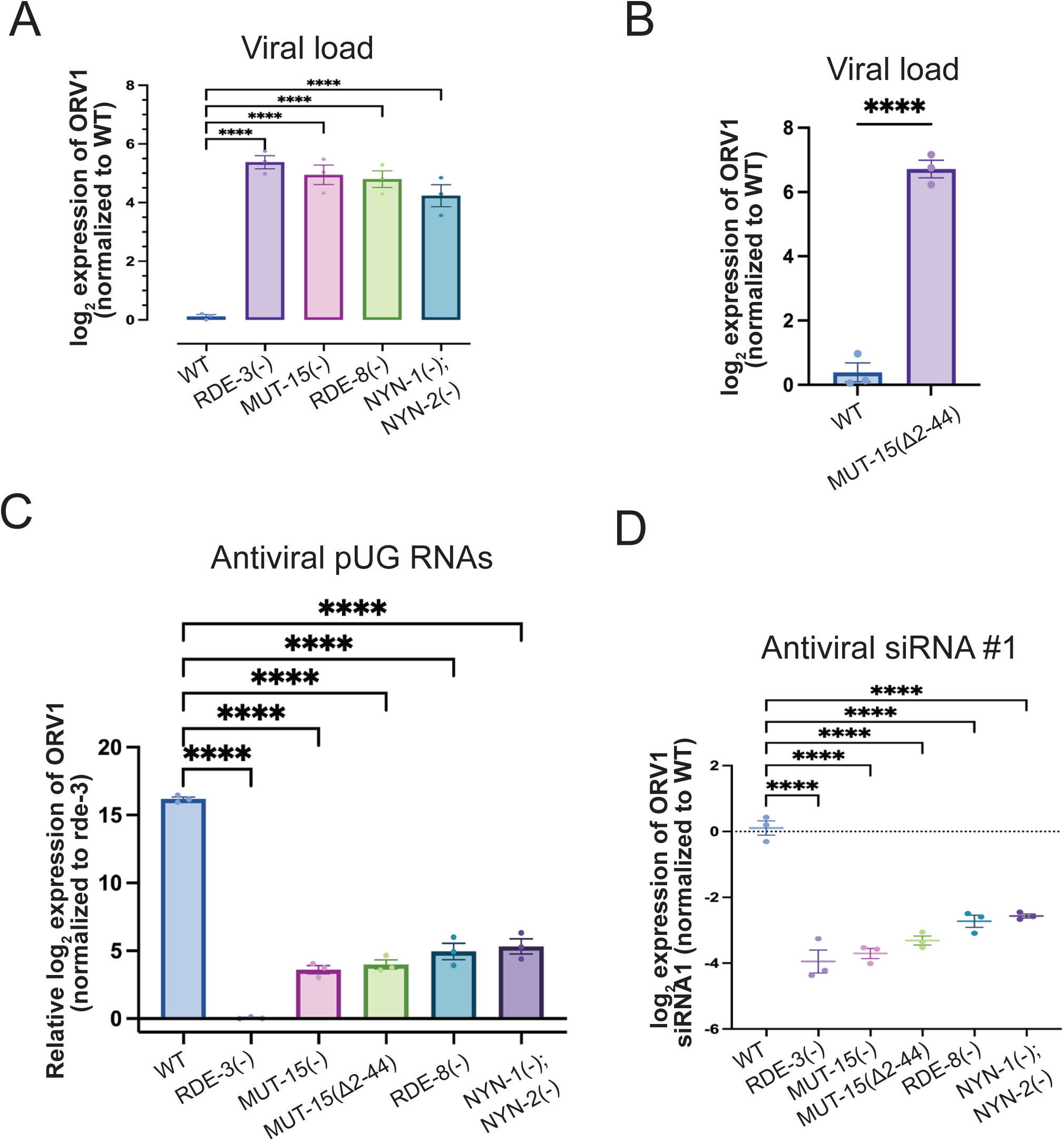
All components of the predicted pUGasome are required for antiviral immunity. **(A-B)** qRT-PCR in **(A)** WT, *rde-3(ne3370)* [RDE-3(-)], *mut-15(tm1358)* [MUT-15(-)], *rde-8(tm2252)* [RDE-8(-)], *nyn-1(tm5004); nyn-2(tm4844)* [NYN-1(-); NYN-2(-)] and **(B)** *mut-15(gg1013)* [MUT-15(Δ2-44)] animals for viral load +/-Orsay infection. Data were normalized to the mRNAs *eft-2* and *cdc-42* and viral loads in WT were defined as zero. Unpaired, one-way ANOVA and unpaired, two-tailed t-test. n=3. **(C)** Animals with genotypes described in **(A)** were infected with Orsay and antiviral pUG RNAs were detected with ORV1 pUG RNA qRT-PCR and normalized to the *gsa-1* mRNA. ORV1 pUG RNA levels in *rde-3(-)* were defined as zero. Unpaired, one-way ANOVA. n=3 **(D)** Taqman assays for antiviral ORV1 siRNA1 on RNA isolated from animals with genotypes described in **(A).** Data were normalized to the control *U18* snRNA and signals in WT were defined as zero. Paired, one-way ANOVA. n=3. ns - not significant,* p<0.05, ** p<0.01, *** p<0.001, **** p<0.0001.

### MUT-16 and RdRP act downstream of the pUGasome

Interestingly, MUT-16(-) animals behaved differently than animals lacking the components of the predicted pUGasome when infected with Orsay virus. While MUT-16(-) animals exhibited elevated viral loads and were defective for synthesizing antiviral siRNAs (Figure 6A/B and Figure S10A/B), similar to animals lacking components of the predicted pUGasome, MUT-16(-) animals retained the ability to produce antiviral pUG RNAs (Figure 6C/D). Thus, MUT-16 is not required for viral pUG RNA synthesis, but is required for antiviral siRNA synthesis and for suppressing viral load.

**Figure 6.**
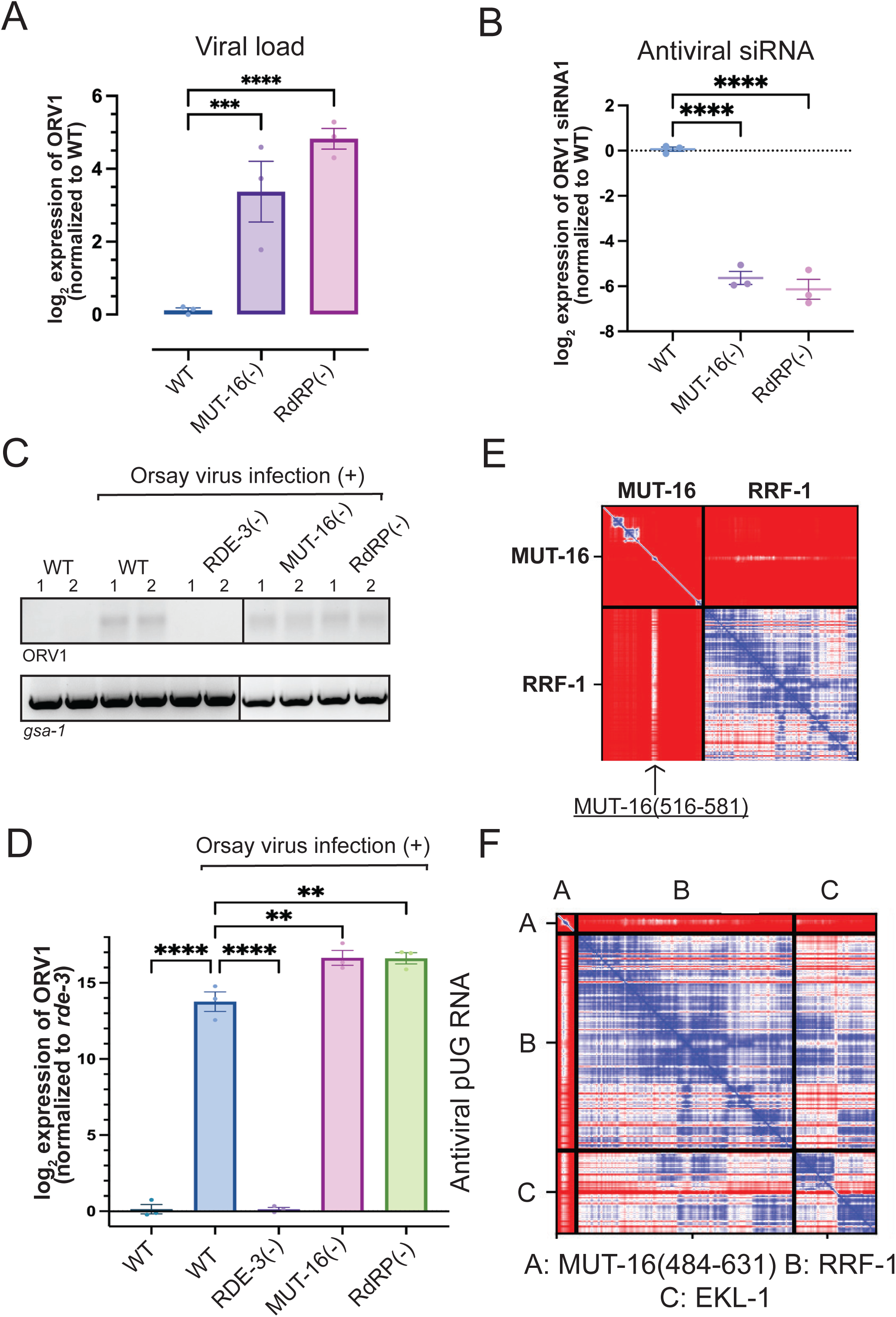
MUT-16 and RdRP function downstream of the predicted pUGasome. **(A)** qRT-PCR using RNA isolated from Orsay infected wild-type [WT], *mut-16(pk710)* [MUT-16(-)], and *rrf-1(pk1417); ego-1(gg685)* [RdRP(-)] animals to assess viral loads. Data are normalized to housekeeping genes *eft-2* and *cdc-42* and signals in WT were defined as one. Unpaired, one-way ANOVA. n=3 **(B)** Taqman assay using RNA isolated from Orsay infected animals of genotypes described in **(A)**. Data were normalized to the control *U18* snRNA and signals in WT were defined as zero. Paired, one-way ANOVA. n=3. **(C)** RT-PCR of +/- Orsay infected animals of genotypes described in **(A)**. RT-PCR of the control *gsa-1* pUG RNA is shown. **(D)** qRT-PCR for ORV1 pUG RNAs in +/- Orsay infected animals of the genotypes described in **(A)**. Data are normalized to the control *gsa-1* pUG RNA. Unpaired, one-way ANOVA. n=3. ns - not significant,* p<0.05, ** p<0.01, *** p<0.001, **** p<0.0001. **(E-F)** Top ranked PAE plot for **(E)** MUT-16-RRF-1 and **(F)** MUT-16(484-631)-RRF-1-EKL-1 Alphafold2 predictions. **(E)** RRF-1 interacting region in MUT-16 (a.a. 516-581) is marked with an arrow. PAE plots are color-coded to show predicted distances between residues: 0 Angstroms (blue) to 30 or greater Angstroms (red).

Why might MUT-16 be needed for antiviral siRNA synthesis, but not antiviral pUG RNA synthesis? MUT-16 co-IPs with the components of the predicted pUGasome as well as with RdRP (Tsai *et al*., 2015; Manage *et al*., 2020). Current models posit that RdRP uses pUGylated RNAs as templates for siRNA synthesis (Shukla *et al*., 2020; Roschdi *et al*., 2022). Thus, MUT-16 could promote antiviral immunity by bringing RdRP into proximity with the pUGasome, thus enabling efficient conversion of pUG RNAs into antiviral siRNAs. Consistent with this idea, we observed that RdRP mutant animals behaved similarly to MUT-16(-) animals following viral infection: they had elevated viral loads and were defective for synthesizing antiviral siRNAs (Figure 6A/B and Figure S10A/B), which is consistent with previous reports (Ashe *et al*., 2013; Guo, Zhang, Wang and Lu, 2013; Le Pen *et al*., 2018; Long, Meng and Lu, 2018). However, RdRP(-) animals retained the ability to produce viral pUG RNAs (Figure 6C/D and Figure S10A/B). The data hint that MUT-16 could promote antiviral immunity by recruiting RdRP into proximity with the pUGasome, thus coupling antiviral pUG synthesis by the pUGasome with antiviral siRNA synthesis by RdRP. The TUDOR-domain protein EKL-1 interacts with the RdRP RRF-1 to promote RdRP-based siRNA biogenesis (Rocheleau *et al*., 2008; Gu *et al*., 2009; Thivierge *et al*., 2011; Phillips *et al*., 2012; Chen *et al*., 2024). We used AlphaFold2-Multimer to ask if RRF-1 and/or EKL-1 might be predicted to interact with each other and/or with MUT-16. Indeed, AlphaFold2-Multimer predicted that RRF-1 and EKL-1 interact with each other (Figure S11A) and with a.a. 516-581 of MUT-16 (Figure 6E/F and Figure S11B-D). PAE plots for the predicted MUT-16-RRF-1-RdRP interaction are shown in Figure 6E/F and Fig S11B-D. A 3D representation of the predicted RdRP-EKL-1-MUT-16(484-632) complex is shown in Figure S11E. The residues in MUT-16 predicted by Alphafold to interact with RRF-1/EKL are distinct from those predicted to interact with MUT-15 and, therefore, the pUGasome. The RdRP-EKL-1-MUT-16 Alphafold2 prediction is experimentally supported by published IP-MS results (Gu *et al*., 2009), and with data showing that a.a. 484-569 of MUT-16 are necessary for RdRP to colocalize with MUT-16 (Uebel *et al*., 2018). Taken together, the data suggest that MUT-16 promotes antiviral immunity by physically linking the pUGasome to RdRP, thereby enabling efficient conversion of pUGylated RNAs into antiviral siRNAs (Figure 7).

**Figure 7.**
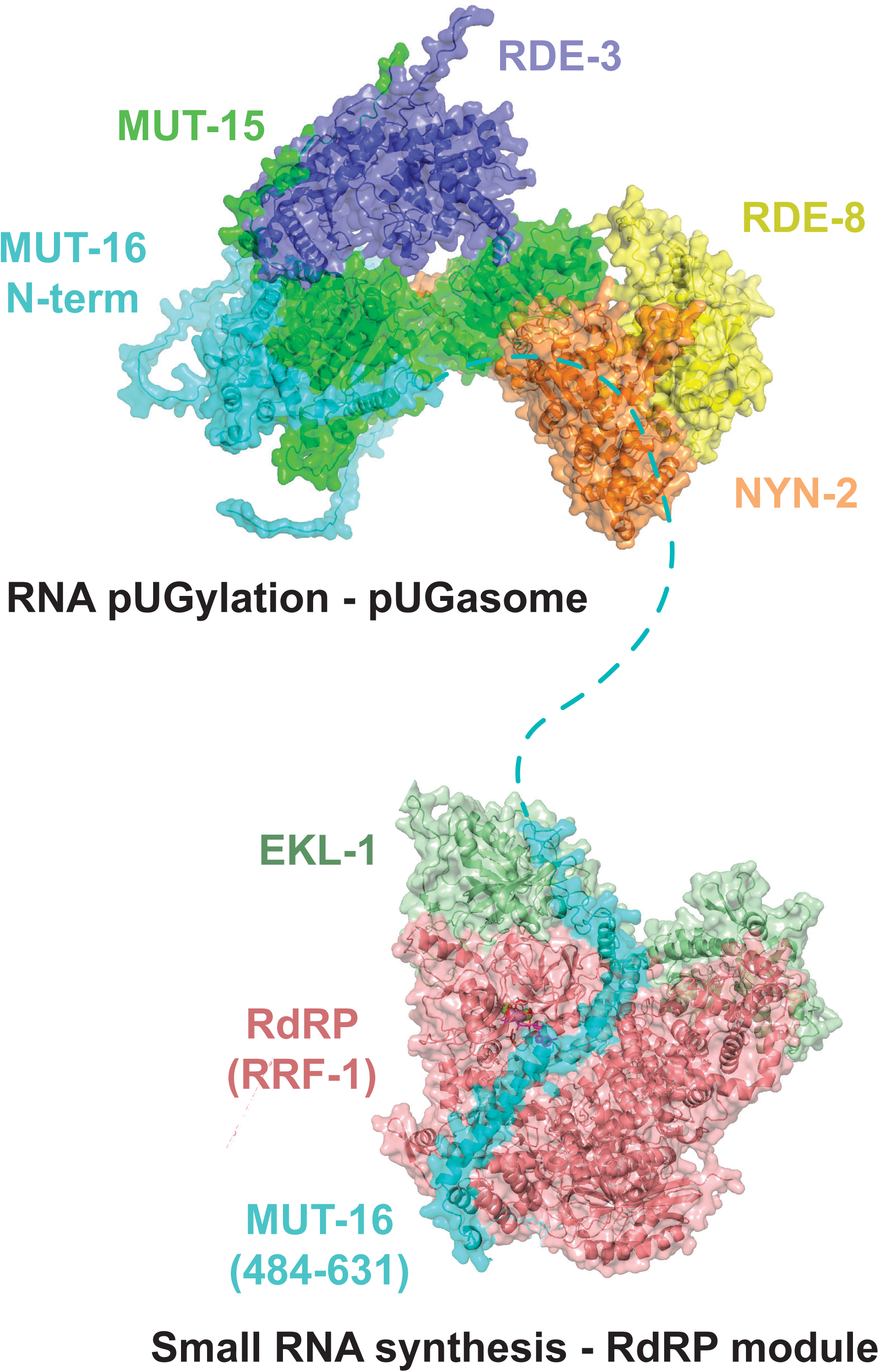
Model for pUG RNA-mediated antiviral immunity. Model of MUT-16 bound to the pUGasome and to RdRP. Predicted components of the pUGasome and the RdRP interacting module are labeled. Structures and interaction surfaces are Alphafold2 predictions.

## Discussion

Here we show that RNA pUGylation limits Orsay virus infection in *C. elegans*. We present evidence supporting the existence of a four protein complex, which we term the pUGasome, that is required for viral RNA pUGylation and antiviral siRNA synthesis. Finally, we show that the pUGasome associates with MUT-16 and this association likely enables antiviral siRNA production by RdRP.

AlphaFold-Multimer simulations predicted that RDE-3, MUT-15, NYN-1/2, and RDE-8 assemble into a pUGasome. Consistent with this model, we find that all four components of the pUGasome are required for viral pUGylation. Biochemical tests of the AlphaFold-Multimer pUGasome prediction support the general architecture of the Alphafold pUGasome prediction: 1) RDE-3 and MUT-15 co-IP *in vivo*; 2) MUT-15(Δ2-44) mutants fail to co-IP with RDE-3, leading to pUGylation defects; 3) RDE-3 co-IPs with RDE-8 and NYN-1 in a MUT-15-dependent manner; and 4) RDE-3 and MUT-15 retain the ability to co-IP in animals lacking MUT-16. We tested one predicted interaction surface of the pUGasome and observed that a.a. 2-44 of MUT-15, which are predicted to mediate contact between MUT-15 and RDE-3, were necessary for MUT-15-RDE-3 interaction *in vivo* and for viral RNA pUGylation. It is not yet clear if the other predicted interaction surfaces within the pUGasome are accurate. Cryo-EM analyses could allow the veracity of other predicted interaction surfaces to be assessed.

The pUGasome model is consistent with existing co-IP data and subcellular colocalization studies of the pUGasome components with one notable exception: MUT-15 is not required for RDE-3 to localize to *Mutator* foci, which are thought to be phase separated germ granules that form in *C. elegans* germ cells (Phillips *et al*., 2012; Uebel *et al*., 2018). This is not the expected result if the MUT-15-RDE-3 interaction is the only mechanism localizing RDE-3 to the pUGasome and *Mutator* foci. It is possible that; 1) RDE-3 interacts with other proteins or RNAs, which localizes RDE-3 independently of MUT-15; or 2) that the logic of pUGylation or pUGasome architecture differ in the soma and germline. Our Alphafold-Multimer screen for RDE-3-interacting proteins identified MUT-15, which was the major focus of this study. The analysis also identified UAF-1, the largest subunit of the U2AF splicing complex, as a candidate RDE-3 interactor. The predicted RDE-3:UAF-1 interaction could be an erroneous prediction by Alphafold or, perhaps, indicative of a surprising connection between the pUGasome and pre-mRNA splicing. Asking if UAF-1 and RDE-3 interact *in vivo* will help distinguish these possibilities.

We speculate that the pUGasome assembles in intestinal cells to promote specificity, processivity, and/or fidelity of RNA pUGylation. During RNAi in *C. elegans*, dsRNA is processed into primary small RNAs by the RNase-III-like enzyme Dicer (Tabara *et al*., 2002). The Argonaute protein RDE-1 binds these primary small RNAs to regulate other cellular RNAs via the recruitment of the endonuclease RDE-8, which cleaves mRNAs targeted by RDE-1 (Tabara *et al*., 1999, 2002; Tsai *et al*., 2015). RDE-3 appends pUG tails to RDE-8 cleaved RNAs and RdRPs then amplify gene silencing responses by using pUGylated RNAs as templates for secondary siRNAs, which engage WAGO proteins to further silence homologous RNAs (Shukla *et al*., 2020). We find that the components of the pUGasome are required for both RNAi and viral RNA pUGylation, suggesting that the logic of viral pUGylation is, at least partly, similar to that of canonical RNAi-based gene silencing in *C. elegans*. The data support a model in which Dicer processed primary small RNAs, derived from viral dsRNAs produced during viral replication, recruit the pUGasome to single-stranded viral RNAs to enable efficient cleavage (RDE-8) and pUGylation (RDE-3) of these RNAs. More specifically, pUGasome assembly may allow the 3’ termini of viral RNAs, generated by RDE-8-based endonucleolytic cleavage, to be brought into close proximity with RDE-3 to enable efficient pUGylation. Indeed, the Alphafold-predicted pUGasome shows a groove of positively-charged surface amino acids connecting the RDE-8 and RDE-3 active sites (Figure S12A/B). Because positively-charged protein surfaces often interact with negatively charged nucleic acids, we speculate that RNA, which is cleaved by RDE-8, is threaded along this positively charged groove to link RNA cleavage with RNA pUGylation (Figure S12A/B). The translocation of RNA along this positively charged groove from RDE-8 to RDE-3 may be facilitated by helicases, such as RDE-12 and MUT-14 (Phillips *et al*., 2014; Shirayama *et al*., 2014; Yang *et al*., 2014). Finally, RDE-3 is remarkable in that it appends alternating uridine and guanosine nucleotides to the 3’ termini of RNAs (Preston *et al*., 2019; Shukla *et al*., 2020). Thus, the RDE-3 active site must alternatively recognize and accommodate either uridine or guanosine nucleotides and these interactions must somehow be coupled to the identity of the last nucleotide added to RNA by RDE-3. It is possible that MUT-15, or MUT-15 in conjunction with the larger pUGasome, could help facilitate the complex structural rearrangements needed for RDE-3 to accomplish this remarkable feat.

MUT-16 is a large (1054 a.a.), low-complexity protein that co-IPs with the subunits of the pUGasome (Phillips *et al*., 2012; Tsai *et al*., 2015; Uebel *et al*., 2018; Manage *et al*., 2020). Alphafold predicts a direct interaction between MUT-16 and MUT-15, which likely explains why MUT-16 co-IPs with pUGasome components. Interestingly, we find that MUT-16 is not needed for antiviral RNA pUGylation. Rather, MUT-16 is needed for antiviral siRNA production. Current models posit that the pUG tails appended to RNAs by RDE-3 interact with RdRP and that this interaction allows RdRP to use pUGylated RNAs as templates to synthesize siRNAs (Shukla *et al*., 2020). MUT-16 also co-IPs with RdRP (Tsai *et al*., 2015; Manage *et al*., 2020). Therefore, we hypothesize that a primary role of MUT-16 during antiviral immunity is to bring RdRP, and likely other factors such as MUT-7, into proximity with the pUGasome so that pUG RNAs can be efficiently converted into the secondary siRNAs, which are loaded onto Argonaute proteins to drive antiviral immunity. Consistent with this model, we find that RdRP is also not needed for pUGylation, but is needed for antiviral siRNA synthesis. Previous studies showing that C-terminus of MUT-16 is needed for MUT-16 to nucleate *Mutator* foci in germ cells and for *in vitro* coacervation (Phillips *et al*., 2012; Uebel *et al*., 2018; Busetto *et al*., 2024). These observations hint that multivalent interactions between the C-termini of MUT-16 could underlie the assembly of *Mutator* foci in germ cells, perhaps via a liquid-liquid phase separation-like process. MUT-16-labeled foci are not observed in the intestinal cells of *C. elegans*, which are a site of antiviral pUG/siRNA biogenesis, suggesting that either MUT-16 oligomerization is a germline-specific function of MUT-16 or that oligomerization also occurs in the soma, but not to a level detectable by light microscopy (Phillips *et al*., 2012; Uebel *et al*., 2018). Asking whether the C-terminus of MUT-16 is needed for MUT-16 to link antiviral pUG synthesis to antiviral siRNA synthesis should distinguish these possibilities.

## Figure Legends

**Figure S1. Auxin-dependent depletion of RDE-3.** Animals of the indicated genotypes were exposed to *dpy-6* RNAi +/- auxin. Animals that failed to respond to *dpy-6* RNAi exhibit normal body sizes, while animals that respond to *dpy-6* RNAi exhibit a Dumpy phenotype.

**Figure S2. Spike-in RNA controls for pUG-Seq.** *In vitro* transcribed gfp(UG)18 RNA was spiked-in to RNA preparations during library preparation. Nanopore sequencing revealed that WT and RDE-3(-) animals had similar levels of spiked-in pUG RNA over three replicates. Unpaired, two-tailed t-test. n=3. ns - not significant,* p<0.05, ** p<0.01, *** p<0.001, **** p<0.0001.

**Figure S3. PAE and 3D representations of Alphafold2 predicted interactions between RDE-3, MUT-15, and MUT-16. (A)** (Top) A predicted 3D structure of the top ranked Alphafold2 prediction for RDE-3 and MUT-15. (Bottom) Five independent PAE plots for the predicted RDE-3-MUT-15 interaction. Rank #1 is also shown in Figure 3. **(B)** Five independent PAE plots for the predicted full-length MUT-16 and MUT-15 interaction. **(C)** (Top) The top PAE plot for the predicted RDE-3-MUT-15-MUT-16(strd) predicted interaction is shown. (Middle) A predicted 3D structure (rotated 180°) of the top ranked prediction of the trimeric complex of RDE-3-MUT-15-MUT-16(strd). Yellow lines connect sections of PAE plots (yellow rectangles) to relevant sections of 3D structures (yellow circles). (Bottom) Five independent PAE plots for RDE-3-MUT-15-MUT-16(strd). PAE plots are color-coded to show predicted distances between residues: 0 Angstroms (blue) to 30 or greater Angstroms (red).

**Figure S4 PAE and 3D representations of Alphafold2 predicted interactions between MUT-15, NYN-1/2, and RDE-8. (A)** Five independent PAE plots for the predicted (top) MUT-15-NYN-1 and (bottom) MUT-15-NYN-2 interactions. **(B) (Top)** Predicted 3D structures and PAE plots for (left) RDE-8 (yellow) and (right) RDE-8 (yellow) and NYN-2 (orange). Red circles and red-colored residues indicate the RDE-8 catalytic site. Corresponding PAE plots are shown. The catalytic site of RDE-8 is labeled in red. PAE plots are measured in Angstroms: 0 (red) to 30 (blue). **(C)** Five independent PAE plots for the predicted (top) RDE-8-NYN-1 and (bottom) MUT-15-NYN-2 interactions. PAE plots are color-coded to show predicted distances between residues: 0 Angstroms (blue) to 30 or greater Angstroms (red).

**Figure S5. Alphafold2 predicts a heteropentamer pUGasome complex.** (Top) The top PAE plots for the predicted MUT-16(strd), RDE-3, MUT-15, RDE-8, and NYN-1 or NYN-2 interaction. (Middle) Predicted 3D structures of pentameric complexes containing NYN-1 (left) and NYN-2 (right). Two images are shown for each prediction, rotated 180°. Inset: RDE-3 catalytic site residues shown in red. (Bottom) Five independent PAE plots for the predicted pentameric complexes with NYN-1 or NYN-2. PAE plots are color-coded to show predicted distances between residues: 0 Angstroms (blue) to 30 or greater Angstroms (red).

**Figure S6. The predicted pUGasome without MUT-16.** As described in Figure S5, except MUT-16(strd) was omitted from the Alphafold2 prediction and only a NYN-1 containing 3D representation is shown. The NYN-2 predicted structure was similar.

**Figure S7. Epitope tags do not disrupt RDE-3, RDE-8, and MUT-15 functions in RNAi. (A)** This panel shows that loss-of-function mutations in the predicted components of the pUGasome are, as expected, defective for dpy-6 RNAi. Animals of the indicated genotypes were exposed to +/- *dpy-6* RNAi. Animals that failed to respond to *dpy-6* RNAi exhibit normal body sizes, while animals that respond to *dpy-6* RNAi exhibit a Dumpy phenotype. **(B)** This panel shows that introduction of epitope tags to the endogenous copies of *rde-8, rde-3, and mut-15* do not disrupt the function of the encoded proteins in *dpy-6* RNAi.

**Figure S8. MUT-15(Δ2–44) is not strongly predicted to interact with RDE-3.** Four of five PAE plots for RDE-3-MUT-15(Δ2–44) and RDE-3-MUT-15(Δ2–44)-MUT-16(strd) do not predict an interaction. PAE plots are color-coded to show predicted distances between residues: 0 Angstroms (blue) to 30 or greater Angstroms (red).

**Figure S9. The predicted pUGasome components are required for antiviral immunity.** The data in this figure show that the predicted components of the pUGasome are required for antiviral immunity as measured by ORV2 levels and ORV1 siRNA #2 quantification. **(A-B)** Genotypes as in Figure 5. **(A)** qRT-PCR for ORV2 RNA levels. Data were normalized to mRNA controls *eft-2* and *cdc-42* and viral loads in WT were defined as zero. Unpaired, one-way ANOVA. n=3. **(B)** Taqman assays for antiviral ORV1 siRNA2. Data were normalized to the control *U18* snRNA and signals in WT were defined as zero. Paired, one-way ANOVA. n=3. ns - not significant,* p<0.05, ** p<0.01, *** p<0.001, **** p<0.0001.

**Figure S10. MUT-16 and RdRP are needed for antiviral siRNA production and antiviral immunity. (A)** qRT-PCR using RNA isolated from Orsay infected wild-type [WT], *mut-16(pk710)* [MUT-16(-)], and *rrf-1(pk1417); ego-1(gg685)* [RdRP(-)] animals to assess ORV2 viral load. Data are normalized to housekeeping genes *eft-2* and *cdc-42* and signals in WT were defined as one. Unpaired, one-way ANOVA. n=3 **(B)** Taqman assay using RNA isolated from Orsay infected animals of genotypes described in **(A)** detecting ORV1 siRNA #2. Data were normalized to the control *U18* snRNA and signals in WT were defined as zero. Paired, one-way ANOVA. n=3. ns - not significant,* p<0.05, ** p<0.01, *** p<0.001, **** p<0.0001.

**Figure S11. Alphafold2 prediction of RdRP-MUT-16 interaction.** Five independent PAE plots for Alphafold2 predicted interactions between **(A)** RRF-1-EKL-1, **(B)** MUT-16-EKL-1, **(C)** MUT-16-RRF-1 and **(D)** MUT-16(484-631)-RRF-1-EKL-1. **(E)** Predicted 3D structure of the top ranked Alphafold prediction for the MUT-16(484-631)-RRF-1-EKL-1 interaction, along with the corresponding PAE plot. PAE plots are color-coded to show predicted distances between residues: 0 Angstroms (blue) to 30 or greater Angstroms (red).

**Figure S12. Surface electrostatic potential of the predicted pUGasome. (A-B)** Electrostatic potential of the Alphafold2 predicted heteropentamer containing **(A)** RDE-3, RDE-8, NYN-2, MUT-15, and MUT-16(strd) and **(B)** RDE-3, RDE-8, NYN-2, and MUT-15. **(A-B)** Two predicted structures, rotated by 180°, are shown for each. **(C)** 3D representation of Alphafold2 prediction that included an RNA in the prediction is shown. Electrostatic potential is indicated with positively-charged residues in red and negatively-charged residues in blue.

## Materials and Methods

### Strains

All *C. elegans* strains were grown at 20 °C, derived from the Bristol N2 strain, and grown on normal growth medium with OP50 bacteria unless otherwise stated. Strains used in this study are listed in Table 1. All the sequences for oligonucleotides, gRNAs, and HR templates are listed in Table 2.

**Table 1.** *C. elegans* strains used in this study.

| Strain name | Genotype | Source and description |
| --- | --- | --- |
| N2 | wild-type | CGC |
| YY1449 | *rde-3(ne3370) I* | *rde-3* deletion - CGC |
| YY1711 | *rde-3(ne3370) I; rde-3(ggSi19) II* | RDE-3 + 2kb upstream and downstream sequence integrated into the LGII MosSCI site (ttTi5605) - Kennedy Lab |
| YY1505 | *rde-3(ne298) I* | G366R catalytic mutation - CGC |
| YY1670 | *rde-3(gg679) I* | R366G reversion to wild-type - Kennedy Lab |
| YY1794 | *rde-3(r459) I* | D189N catalytic mutation - CGC |
| YY1868 | *rde-3(gg693) I; ieSi57 [eft-3p::TIR1::mRuby::unc-54 3'UTR + Cbr-unc-119(+)] II* | pan-soma TIR1 expression with GFP::AID::RDE-3 - Kennedy Lab |
| YY2671 | *rde-3(gg693) I; ieSi57 [myo-2p::TIR1::mRuby::unc-54 3'UTR + Cbr-unc-119(+)] II* | muscle TIR1 expression with GFP::AID::RDE-3 - Kennedy Lab |
| YY2672 | *rde-3(gg693) I; ieSi57 [ges-1p::TIR1::mRuby::unc-54 3'UTR + Cbr-unc-119(+)] II* | intestinal TIR1 expression with GFP::AID::RDE-3 - Kennedy Lab |
| YY2454 | *rde-3(gg636); mut-15(gg1012) I* | 3X FLAG::RDE-3 and MUT-15::3X HA - Kennedy Lab |
| YY2455 | *rde-3(gg636); mut-15(gg1013) I* | 3X FLAG::RDE-3 and 3X |
| | | HA(Δ2-44)::MUT-15 - Kennedy Lab |
| YY2499 | *rde-3(gg636); mut-15(gg1050) I rde-8(gg1051) IV* | 3X FLAG::RDE-3, MUT-15::3X HA, and 3X V5::RDE-8 - Kennedy Lab |
| YY2501 | *rde-3(gg636); mut-15(gg1050) I nyn-1(gg1052) IV* | 3X FLAG::RDE-3, MUT-15::3X HA, and 3X V5::NYN-1 - Kennedy Lab |
| YY2502 | *rde-3(gg636); mut-15(gg1050); nyn-2(1053) I* | 3X FLAG::RDE-3, MUT-15::3X HA, and 3X V5::NYN-2 - Kennedy Lab |
| YY2513 | *rde-3(gg636); mut-15(gg1067) I nyn-1(gg1052) IV* | 3X FLAG::RDE-3, MUT-15::3X HA with R159STOP mutation, and 3X V5::NYN-1 - Kennedy Lab |
| YY2522 | *rde-3(gg636); mut-15(gg1067); nyn-2(1053) I* | 3X FLAG::RDE-3, MUT-15::3X HA with R159STOP mutation, and 3X V5::NYN-2 - Kennedy Lab |
| YY2555 | *rde-3(gg636); mut-15(gg1067) I rde-8(gg1051) IV* | 3X FLAG::RDE-3, MUT-15::3X HA with R159STOP mutation, and 3X V5::RDE-8 - Kennedy Lab |
| YY1689 | *ego-1(gg685) rrf-1(pk1417) I* | null alleles for both host RdRPs, which are in an operon - Kennedy Lab |
| GR1747 | *mut-15(tm1358) I* | Deletion of MUT-15 - CGC |
| YY1803 | *nyn-1(tm5004) I; nyn-2(tm4844) rde-8(tm2252) IV* | Deletion of RDE-8, NYN-1, and NYN-2 - Mello Lab |
| YY1740 | *nyn-1(tm5004) I; nyn-2(tm4844) IV* | Deletion of NYN-1 and NYN-2 - Mello Lab |
| YY1739 | *rde-8(tm2252) IV* | Deletion of RDE-8 - Mello Lab |
| NL1810 | *mut-16(pk710) I* | Null allele of MUT-16 - CGC |

**Table 2.**
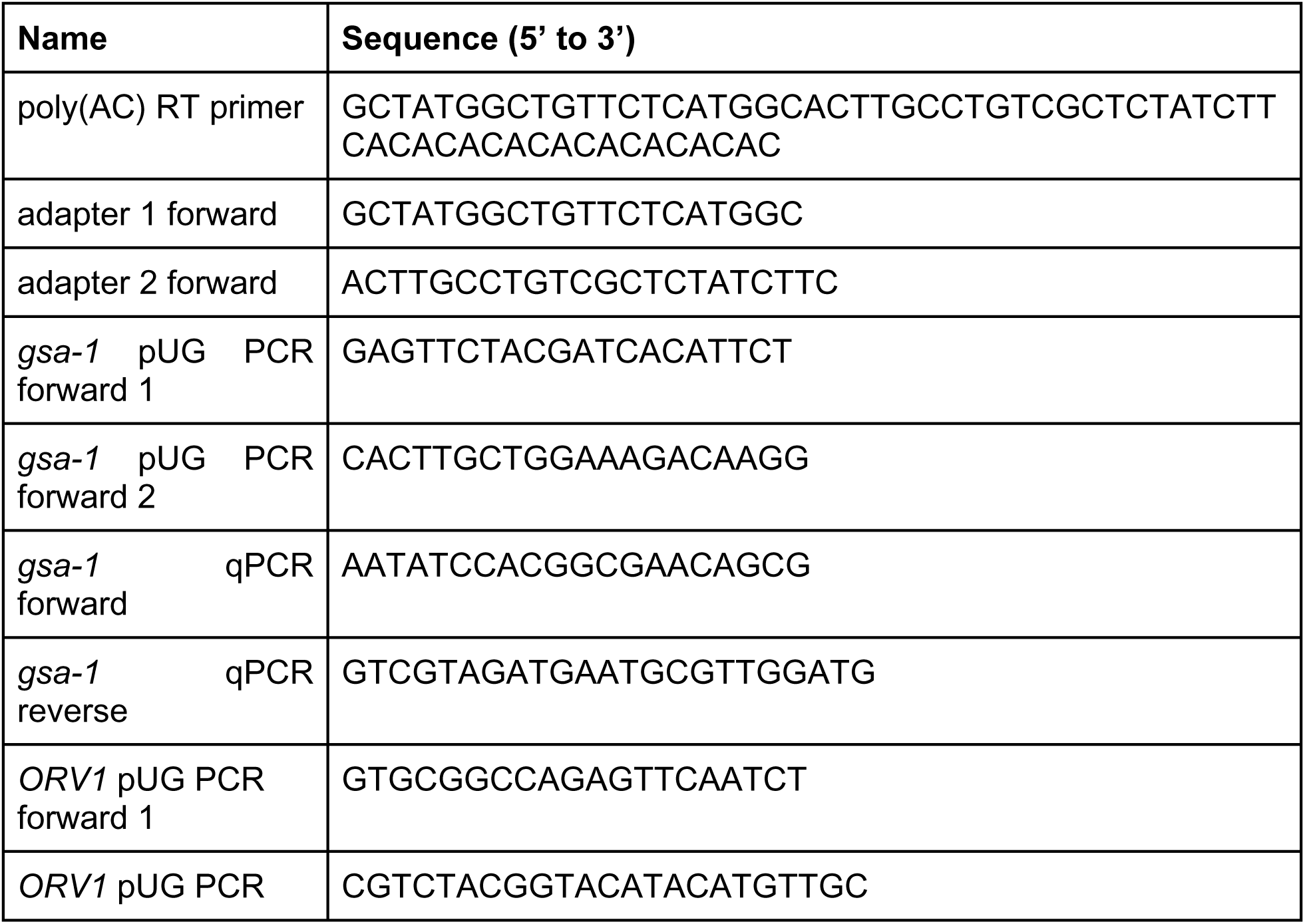
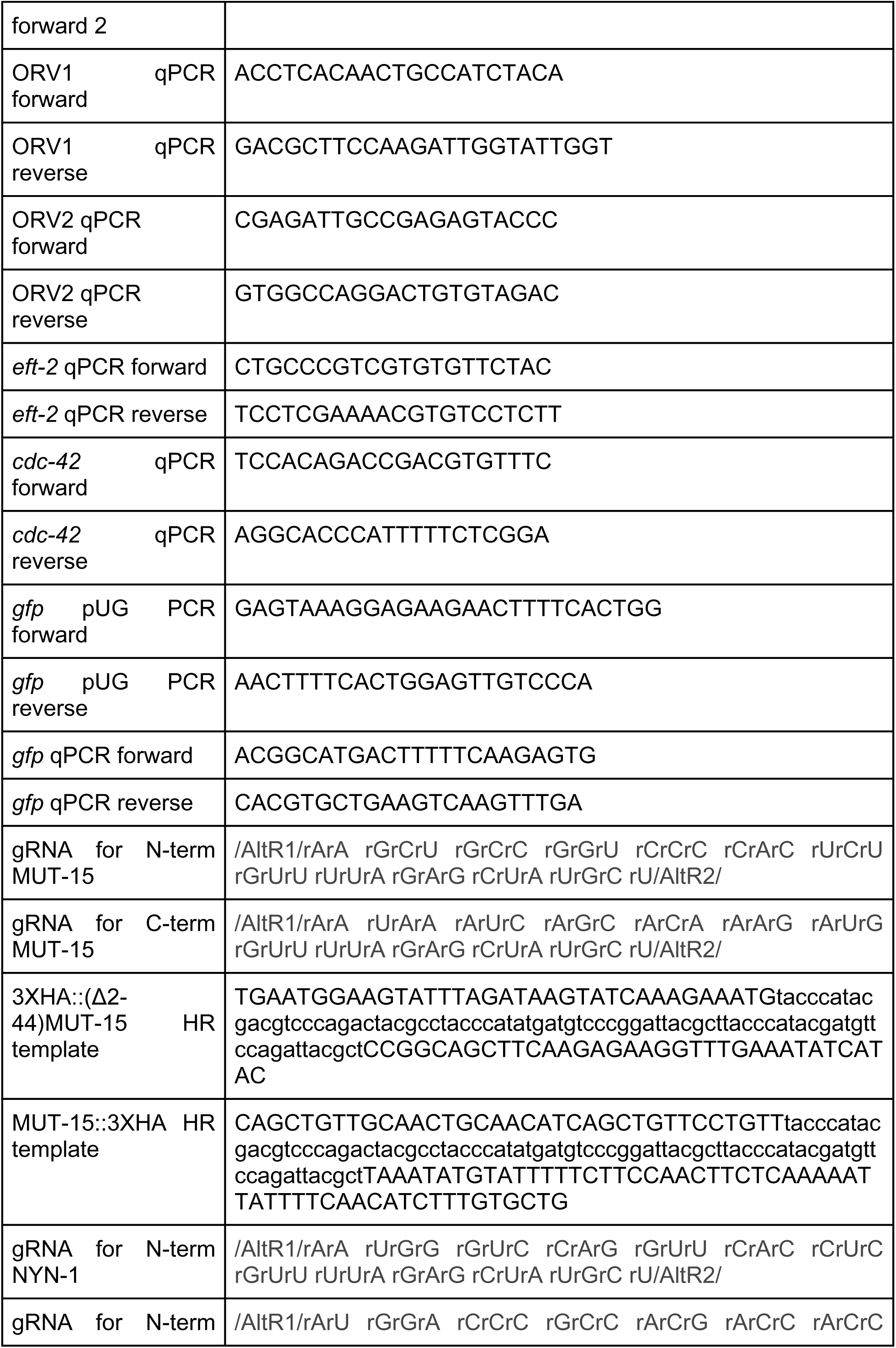
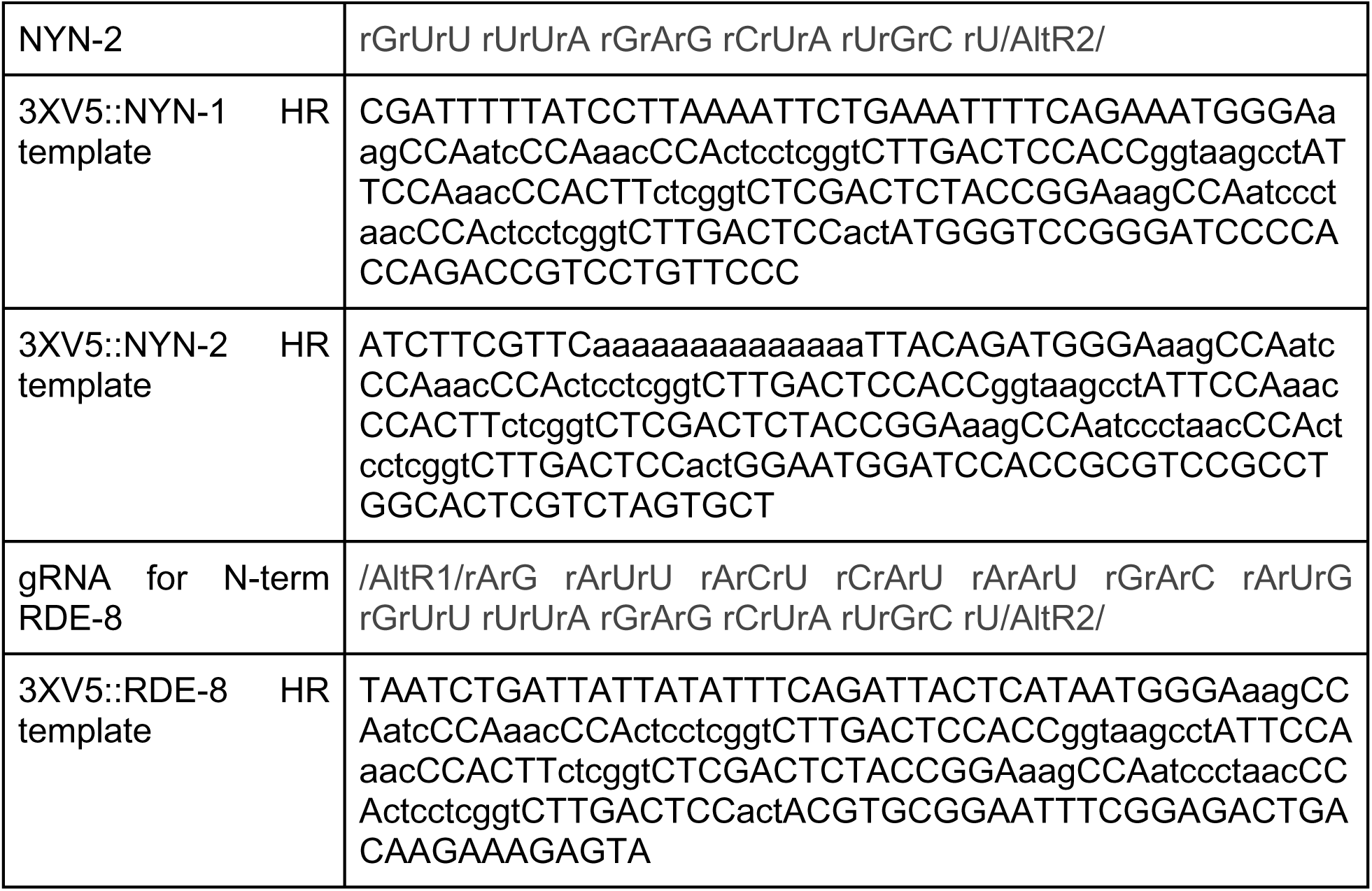
Oligonucleotides, gRNAs, and HR templates used in this study.

### RNA Extraction

Animals were collected in TRIzol Reagent and freeze-thawed at least three times. RNA was extracted with Isoamyl-alcohol and chloroform twice or with RNA clean and concentrator with on-column DNAse digestion (Zymo Research, R1014). RNA was quantified using a Nanodrop2000 to determine concentration.

### CRISPR-Cas9

Gene editing preparation in C. elegans was performed as previously described (Dokshin *et al*., 2018). Homology Repair templates were ordered from IDT as Alt-R HR oligos, if possible. Custom sgRNAs were designed for genes of interest using Benchling. The designed sgRNAs were validated using IDTs custom gRNA tool. CRISPR-Cas9 complex was assembled as previously described (Paix *et al*., 2015). Adult animals were injected in the germline and F1s were phenotyped for the co-injection marker of an extrachromosomal array (*rol-6*). F2s were singled and non-roller animals were maintained and genotyped. All the edited animals were validated with Sanger sequencing and outcrossed at least once.

### pUG-Seq Library Preparation

Spike-in RNA (*gfp(UG)18*) was synthesized using the Megascript T7 kit (Invitrogen, AM1334) and purified using the RNA Clean Concentrator (Zymo Research, R1014). RNAseOUT (Invitrogen, 10777019) was added with RNA to prevent unwanted degradation. A combination of 10 ug of total RNA and 100 pg of spike-in RNA was depleted of rRNA using a previously established RNAse-H (NEB, M0297S) based method (Gallucci et al). The rRNA-depleted RNA was purified using RNA clean and concentrator (Zymo Research, R1014). The eluted RNA was used for reverse transcription with Maximus H Minus Reverse transcriptase (Thermo Fisher, EP0742), Strand Switching Primer II (SSP II), and a poly(AC) RT primer according to the Oxford Nanopore cDNA-PCR barcoding protocol. To remove unwanted fragments from the sequencing library, the newly generated cDNA was initially amplified with 2X Longamp Taq Polymerase (NEB, M0287L) for 8 cycles to generate dsDNA. DASH was performed to remove any *yrn-1* and RT primer-SSP II artifacts. Briefly, Cas9–gRNA complex (IDT) was prepared exactly as previously described (Gu *et al*., 2016; Mahat *et al*., 2024) and pre-incubated for 15 min at 25 °C. The incubated complex was used to digest the cleaned dsDNA library by incubating for 1 hour at 37 °C. The digested samples were purified using a select-a-size concentrator (Zymo Research, D4080) to deplete fragments <300 nts and eluted with nuclease-free water. The final library was analyzed with Tapestation (Agilent) before proceeding to sequencing.

### Nanopore Sequencing

The library was pooled and prepared according to Nanopore protocol. Briefly, the pooled library was loaded into MinION r10 flow cells on a MinION Mk1C machine (Oxford Nanopore). Sequencing was set to run for at least 24 hours, barcoding was turned on with no trimming, and quality score filter of 9. Guppy high accuracy base calling model was used.

### Data Preprocessing and Transcript Quantification

*Fastq* files that passed the quality score filter were in the fastq_pass folder. The *fastq* files were merged using *cat *.fastq*. Next, *pychopper* was used to identify and orient full-length transcripts. The PCS111_primers.fas was modified as following:

*>VNP*

*TTGCCTGTCGCTCTATCTTC*

*>SSP*

*TCTGTTGGTGCTGATATTGCTTT*

Pychopper was used with the following settings: *-m edlib -p -k PCS111*. The output fastq files were then merged and filtered for TG repeats using the following:

*grep -B1 -A2 --no-group-separator -E. GTGTGTGTGT.{1\}*

Then, the filtered *fastq* files were then used to map to the transcriptome, spike-in, and the Orsay virus using *minimap2 -uf k14*. The resulting sam files were converted to bam files that were indexed and sorted using *samtools*. *Salmon* was used to generate a *quant.sf* file. The quantification file was imported into R using *tx2import*. The libraries were normalized to RPM using the spike-in *gfp* pUG RNAs to normalize the libraries for any technical variations from the library generation. To analyze visually, reads that were generated from mapping the *fastq* files to the genome were loaded onto the integrated genome viewer or IGV (Broad institute). Sam files were processed using *pysam* to analyze the soft clipped sequences that correspond to the pUG tails as well as identifying the location of pUG tail sites. Frequency of pUG tails across the viral genome (binwidth = 50) were generated using *ggplot2* in R.

### Computational Predictions

All protein sequences were identified from the Uniprot database. To identify conserved germline proteins, we first started with a dataset of germline-specific transcripts identified through SAGE (Wang *et al*., 2009), which was a total of 1063 genes. We then used BioMart on the Wormbase ParaSite to select from the 1063 germline-specific genes that are conserved in humans. We then selected one hundred conserved germline-specific genes without overlapping paralogs when possible. In addition, we added *Mutator* proteins identified in MUT-16 IP-MS (Phillips *et al*., 2012, 2014; Manage *et al*., 2020). The protein sequences of the final list of genes to test for RDE-3 interaction were extracted from Uniprot; if there were multiple isoforms then the longest isoform was chosen for the initial predictions. Alphafold2 was used to perform protein:protein interactions with the following parameters: 5 models and 5 cycles (paste the code). To analyze the outputs, we used a python script to summarize the predictions (Lim *et al*., 2023). To generate the graphs of the *in silico* predictions, we used the avg_n_models per contact and average ipTM scores with *ggplot2* in R. To create the protein structure prediction figures, the top ranked PDB file was loaded onto Pymol. The pLDDT scores were used to identify strongly and poorly predicted regions of the protein:protein interaction.

### Immunoprecipitation

Animals were mixed with 1X RIP Buffer (20 mM Tris-HCl pH 7.5, 200 mM NaCl, 2.5 mM MgCl2, 10% glycerol, 0.5% NP-40, 80 U ml−1 RNaseOUT, 1 mM dithiothreitol (DTT) and protease inhibitor cocktail without EDTA) and frozen dropwise with liquid nitrogen to generate frozen worm balls. The frozen worm balls were ground with a mortar and pestle, before resuspending in the same buffer as collection. The ground lysate was clarified by a centrifuge running at 15000 x g for 10 mins in 4 °C. The supernatant was resuspended into a new tube. The protein lysate was measured for abundance with a BCA assay (Thermo Fisher, 23225). For co-immunoprecipitation assays, anti-FLAG M2 magnetic beads (Thermo Fisher, A36797) were used to immunoprecipitate 3X FLAG::RDE-3 from ∼300-400 ug of protein lysate. Co-immunoprecipitated proteins were eluted with 0.1M glycine pH 3.5 and 1M Tris-HCl pH 8.0 for neutralization; and mixed with Laemmli buffer prior to boiling for loading. Roughly the same amount of proteins is loaded for input and IP unless otherwise stated into a 10% mini-Protean TGX precast SDS-PAGE gels (Biorad, 4561033). The gels were semi-dry transferred using 0.2 uM nitrocellulose Trans-blot Turbo (Biorad, 1704157) and the Trans-blot Turbo Transfer System (Biorad). The membranes were blocked for 1 hour with the Intercept (TBS) blocking buffer (Licor, 927-60003) at room temperature on a shaker. Primary antibodies were incubated overnight at 4 °C on a shaker, then washed four times with 1X TBS-T. The following primary antibodies were used: mouse anti-FLAG (Sigma, F3165, diluted 1:2000), rabbit anti-HA (Abcam, C29F4, diluted 1:2000), rabbit anti-V5 (Cell Signaling Tech, D3H8Q, diluted 1:1000). Secondary antibodies were incubated for at least 1 hour at room temperature, then washed four times with 1X TBS-T prior to imaging on a Licor Odyssey Fx. The following secondary antibodies were used: anti-rabbit IR-680 (Licor, 1:10000) and anti-mouse IR-800 (Licor, 1:10000).

### Orsay Virus Filtrate Preparation and Infection Assays

Orsay virus filtrate was prepared according to previous publications (Sowa *et al*., 2020). Briefly, *rde-1(ne219)* animals infected with Orsay virus were grown for several generations before collecting starved animals with an M9 buffer. The animals were incubated in the buffer for at least 30 mins. Then, the supernatant was collected and isolated from the pellet containing the starved animals. The virus-containing supernatant was plunged through a 22 gauge needle, then isolated into 1 mL virus aliquots to avoid freeze/thaw cycles. The virus aliquots were diluted 1:1000 with 10X OP50 to seed infection plates. Animals were infected by egg prepping onto the Orsay virus-containing plates. Animals were collected at adulthood about 3-4 days post-infection. To infect animals with auxin-mediated depletion of RDE-3 in various tissues, 1mM auxin plates were seeded with virus and various strains were egg prepped onto the infection plates for 3 days prior to collection.

### qRT-PCR Assay

Total RNA were reverse transcribed with the Superscript IV kit (Thermo Fisher) according to the manufacturer’s protocol. Random hexamer was used for quantifying total viral RNA levels from 1 ug of total RNA. A poly(AC)12 with 5’ template sequence encoding an amplification sequence was used to detect pUG RNAs from 5 ug of total RNA. An oligo(dT)20 was used to reverse transcribe polyadenylated mRNAs from 1 ug of total RNA. 1:10 diluted cDNA was mixed with SYBR Green Master Mix (Biorad) and primers for qPCR according to manufacturer’s recommendation in a CFX Connect machine (Biorad). The CT values were used to calculate relative expression levels using the −2^ΔΔ^ method. To quantify viral RNA levels across samples, primers targeting *ORV RNA1 (ORV1)*, *ORV RNA2 (ORV2)*, *cdc-42*, and *eft-2* were used. To quantify pUG RNAs, *gsa-1* primers were used as control.

### Taqman Assay

1 ug of Trizol-extracted RNA from infected animals (see above) was used for Taqman assays. Small RNAs were reverse transcribed into cDNA using the Taqman MicroRNA Reverse Transcription Kit (Applied Biosystems, 4366596). *U18*, antiviral *siRNA1*, and antiviral *siRNA2* small RNAs were then quantified by qRT-PCR using TaqMan Universal Master Mix II, no UNG (Applied Biosystems, 4440040) and custom TaqMan small RNA assays from Applied Biosystems (assay IDs: *U18:* 001764, *Orsay virus RNA1* siRNA1: CTU63TC, *Orsay virus RNA1* siRNA2: CTWCXC9). qRT-PCRs were performed using the CFX Connect machine (Bio-Rad) and semi-skirted PCR plates (Bio-Rad, 2239441).

### Feeding RNAi Assay

RNAi clones were from the Ahringer Library (Source Biosciences). RNAi clones were grown on the appropriate antibiotic. Before seeding, IPTG was added to the RNAi clone and incubated on a shaker at 37 °C for 1 hour. The incubated RNAi clones were 2X concentrated, then seeded onto IPTG plates to express dsRNA. Animals were egg prepped onto the appropriate RNAi plates. *L4440* was used as a negative control. For *dpy-6* RNAi, after 3 days of incubation the plates were inspected and counted for a detectable dumpy phenotype with a light microscope (Nikon). The counts were analyzed and displayed using Graphpad (Prism).

### Auxin-mediated degradation

Animals containing degron tagged RDE-3 with TIR1 expression in either the soma, intestine, or muscle were egg prepped onto 1 mM auxin plates. Animals were grown for 4 days before collection for imaging. Animals were washed off plates using M9 + Triton X-100 buffer, which were washed with M9 buffer. 0.1% sodium azide was added before mounting onto slides with freshly made 2% agarose pad for imaging. For RNAi assay with auxin, 1 mM auxin plates containing IPTG and seeded with *L4440* or *dpy-6* RNAi clones from the Ahringer Library (Source Bioscience). Animals were egg prepped onto the corresponding auxin + RNAi plates and animals were collected after 4 days of incubation. All images were taken using Axio Observer.Z1 fluorescent microscope (Zeiss) with the Plan-Apochromat 20x/0.8 M27 objective.

### Quantifications and Statistical Analysis

All statistics were performed using Graphpad (Prism). The p-values and comparisons relevant to the text are shown in the figures. All descriptions of the statistical analysis are described in the figure legends.

## Author Contributions

D.D.L: Conceptualization, Data Curation, Investigation, Formal analysis, Methodology, Validation, Software, Writing–original draft. A.S: Conceptualization, Investigation, Validation, Resources. S.G.K: Conceptualization, Supervision, Funding acquisition, Writing–original draft, Writing–review and editing.

## Acknowledgements

We thank all the members of the Kennedy Lab for fruitful discussions; Longwen Zhao for technical assistance with genome editing; Fischer/Kim Lab for helpful suggestions; Hope Merens and Churchman Lab for Nanopore assistance; Harvard’s O2 cluster for setting up colabfold to run predictions; Emily Troemel and Troemel Lab for Orsay virus reagents and helpful discussions; Mello Lab for worm strains; Sam Butcher, Bradley Klemm, and Traci Hall on discussions about the pUGasome. We thank Harvard University, Harvard Medical School, and the Department of Genetics for support following the government’s termination of funding to the Kennedy lab.

